# Transcranial magnetic stimulation of the occipital cortex interferes with foot movements in acquired but not congenitally blind individuals

**DOI:** 10.1101/2021.08.03.454870

**Authors:** Tsuyoshi Ikegami, Gen Miura, Masaya Hirashima, Eiichi Naito, Satoshi Hirose

**Author notes:** Correspondence should be addressed to Tsuyoshi Ikegami.

## Abstract

Research has shown that the occipital cortex can be reorganized and repurposed for nonvisual perception and cognitive functions following visual loss in blind individuals. However, no studies have directly examined the involvement of the visual cortex in motor function. Here, we show that a rhythmic foot movement performed by acquired blind individuals can be disrupted by transcranial magnetic stimulation (TMS) to their primary and secondary visual cortex (V1/V2). This disruptive effect was absent for congenitally blind or sighted individuals. As a control, TMS to the neck muscles of acquired blind individuals confirms that this disruptive effect was not caused solely by the sensory (tactile/auditory) sensations accompanying the stimulation. Our results provide the direct evidence that functional repurposing of the human visual cortex is not restricted to perception and cognitive functions but also extends to motor function. Moreover, our findings suggest the nessecity of the visual experience before visual loss for the functional reorganization of the visual cortex for motor function.

**Significance statement:** The brain can adapt remarkably after sensory loss. One striking example is that after visual loss, the visual cortex—deprived of visual input—is often reused for nonvisual perception and higher cognition. Here, we show that it can also support motor control. Using transcranial magnetic stimulation, we transiently disrupted processing in the visual cortex while participants performed rhythmic foot movements. This disruption impaired performance in people who became blind after birth, but not in sighted individuals or those blind from birth. These findings demonstrate that the visual cortex can take on a motor function after vision is lost, but only with prior visual experience.

## Introduction

Sensory loss can lead to dramatic plasticity of the cerebral cortex (*1, 2*). Accumulating evidence indicates cross-modal plasticity following visual loss, where the visual cortex is reorganized to participate in the remaining sensory modalities. For example, the visual cortex of both congenitally and acquired blind individuals responds to auditory (*3–6*) and tactile stimuli (*7–11*). This cross-modal plasticity has been regarded as the neural underpinnings driving blind individuals’ superior ability over sighted individuals in nonvisual perception including sound localization and tactile spatial discrimination (*12–15*). Furthermore, researchers have suggested that the visual cortex can also be reorganized to contribute to higher cognitive processes such as verbal memory, language processing, and mathematical processing (*16–20*). Thus, the visual cortex of blind individuals takes a more prominent role in nonvisual perception and cognitive functions compared with that of sighted individuals.

Despite this understanding, few studies have examined whether the visual cortex can be reorganized to contribute to sensorimotor control. Previous studies have reported that the primary visual cortex (V1) of blind individuals can be recruited for cognitive tasks involving motor output, such as braille reading tasks (*7–9*). However, the recruitment of V1 has only been attributed to their nonvisual perception or cognitive function, but not to motor function. To our knowledge, a single study has suggested that the visual cortex may be reorganized to participate in sensorimotor control involving the spared sensory modalities, with blind opossums being superior to sighted ones in somatosensory-based motor control during ladder-rung walking (*21*). However, direct evidence is still lacking in humans and other animals. To directly investigate whether the visual cortex can be reorganized for sensorimotor control, we applied transcranial magnetic stimulation (TMS) to the occipital cortex, including the primary and secondary visual cortex (V1/V2), of blind participants during a rhythmic foot movement.

There is another debate on differences in the cross-modal plasticity between acquired blinds who lost their vision after birth and congenitally blind individuals who never have experienced vision (*2, 22, 23*). Studies have suggested that the neuronal structure (*24–26*) and activation patterns (*9, 27–29*) of the visual cortex as well as the behavioral performance (*30–32*) differ between acquired and congenitally blinds. Particularly for sensorimotor control, we expected that the reorganization process for recruiting the visual cortex may differ between acquired blinds with experience in visuomotor control and congenitally blinds without it. Therefore, we examined the above question separately for congenital and acquired blind participants.

We observed that TMS to the occipital cortex increased the variability of the foot movement in acquired blind participants but not in congenitally blind participants or sighted controls. Furthermore, we demonstrated that the disruptive effect was not observed when we applied TMS to the blinds’ neck muscle of acquired blind individuals, which induced tactile/auditory sensations similar to those induced by stimulation to the occipital cortex without affecting neural processing. This evidence suggests that the tactile/auditory sensations accompanying TMS alone were not sufficient to cause this disruptive effect. Our findings suggest that a reorganization of the visual cortex in blind individuals contributes to sensorimotor control, requiring some visual experiperience before they lose vision.

## Results

### Experiment-1

Twelve acquired blind individuals, 12 congenitally blind individuals, and 12 sighted individuals participated in the experiment (see Table 1 for participant characteristics). Blindfolded participants in all groups performed rhythmic movements of alternating dorsiflexion and plantar flexion of the feet (Fig. 1A) without any auditory cue. The participants were instructed to maintain their movement frequency (1 Hz) as constant as possible. We chose this task to minimize the influence of differences in visual experience on task performance. In daily life, sighted individuals often control foot movements without visual input, unlike upper limb movements. Moreover, the task requirement―rhythmic dorsiflexion–plantarflexion―can be easily understood without reference to the visual coordinate system. Therefore, task performance was expected to be minimally affected by the amount of the visual experience, allowing a fair comparison among the groups.

**Fig. 1.**
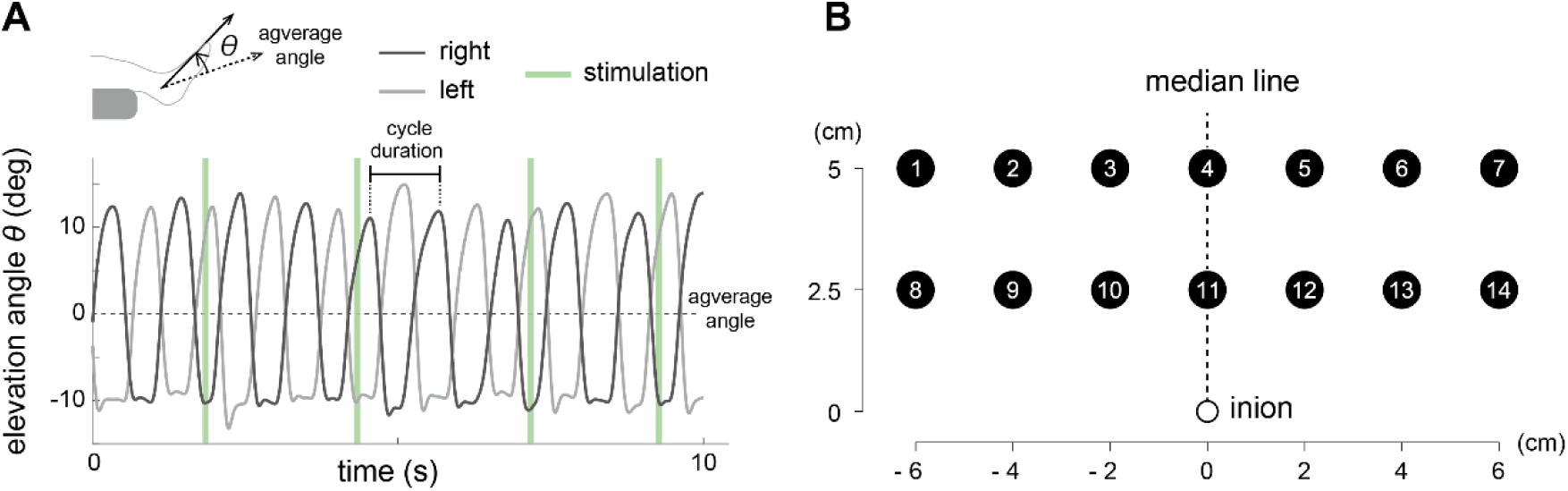
The movement task and stimulation sites: A) The rhythmic foot movement of a representative participant for a 10-s period is shown by the elevation angle, θ of right foot (dark gray line) and left foot (light gray line). For each foot in each trial, θ is defined as the angle relative to the average angle of the trial (θ = 0, dashed line). Each light-green vertical line indicates the timing of a single transcranial magnetic stimulation (TMS) pulse delivered at random. The cycle duration was defined as the time between the adjacent dorsiflexion peaks. The SD of the cycle durations was calculated for each foot in each trial (see **Methods**). B) Fourteen stimulation sites arranged in a 2 × 7 grid were identified for each participant using the inion and the median line as reference points (see **Methods**). Stimulation site #11 corresponds to putative primary and secondary visual cortex (pV1/V2).

**Table 1.** Characteristics of the participants in Experiment 1.

| Participant # | Group | Age (y) | Sex | Onset of blindness (y) | Cause of Blindness | Light sensitivity at present | TMS intensity (%) |
| --- | --- | --- | --- | --- | --- | --- | --- |
| 1 | ABL<sup>A</sup> | 26 | m | 2 | Retinoblastoma | none | 80 |
| 2 | ABL<sup>A</sup> | 27 | m | 14 | retinal choroidal degeneration | yes | 75 |
| 3 | ABL<sup>A</sup> | 39 | m | 20 | morning glory anomaly | none | 85 |
| 4 | ABL<sup>A</sup> | 28 | m | 23 | Uveitis | yes | 90 |
| 5 | ABL<sup>A</sup> | 20 | m | 2 | Retinoblastoma | none | 75 |
| 6 | ABL<sup>A</sup> | 40 | m | 25 | retinitis pigmentosa | yes | 85 |
| 7 | ABL<sup>NA</sup> | 25 | m | 14 | chorioretinal atrophy | yes | 70 |
| 8 | ABL<sup>NA</sup> | 25 | m | 5 | Glaucoma | none | 65 |
| 9 | ABL<sup>NA</sup> | 42 | m | 17 | uveitis + glaucoma | occasional | 70 |
| 10 | ABL<sup>NA</sup> | 19 | m | 2 | behcet's disease | none | 100 |
| 11 | ABL<sup>NA</sup> | 20 | m | 3 | retinal cell death | none | 75 |
| 12 | ABL<sup>NA</sup> | 19 | m | 15 | Glaucoma | none | 55 |
| 13 | CBL | 29 | m | 0 | unknown | none | 80 |
| 14 | CBL | 40 | m | 0 | cataract | none | 95 |
| 15 | CBL | 36 | m | 0 | anophthalmia + persistent fetal vasculature | none | 60 |
| 16 | CBL | 32 | m | 0 | amaurosis | yes | 75 |
| 17 | CBL | 36 | f | 0 | crystalline lens tissue hyperplasia + retinal detachment | yes | 80 |
| 18 | CBL | 35 | m | 0 | Leber congenital amaurosis | none | 65 |
| 19 | CBL | 48 | m | 0 | retinoblastoma | none | 90 |
| 20 | CBL | 43 | m | 0 | unknown | yes | 80 |
| 21 | CBL | 42 | m | 0 | retinoblastoma | none | 55 |
| 22 | CBL | 48 | f | 0 | Retinopathy of prematurity | yes | 80 |
| 23 | CBL | 48 | m | 0 | Corneal staphyloma | none | 80 |
| 24 | CBL | 49 | f | 0 | Retinopathy of prematurity | none | 40 |
| 25 | SI | 22 | m | - | - | - | 85 |
| 26 | SI | 22 | m | - | - | - | 85 |
| 27 | SI | 23 | m | - | - | - | 95 |
| 28 | SI | 22 | m | - | - | - | 90 |
| 29 | SI | 34 | m | - | - | - | 70 |
| 30 | SI | 35 | m | - | - | - | 85 |
| 31 | SI | 40 | m | - | - | - | 80 |
| 32 | SI | 26 | m | - | - | - | 75 |
| 33 | SI | 23 | m | - | - | - | 70 |
| 34 | SI | 27 | m | - | - | - | 100 |
| 35 | SI | 20 | m | - | - | - | 95 |
| 36 | SI | 39 | m | - | - | - | 85 |

*ABL, acquired blind; CBL, congenitally blind; SI, sighted; y, year; m, male. Superscripts A and NA indicate athlete and nonathlete blind participants, respectively.*

During the movement, 20 single TMS pulses were applied to one of 14 stimulation sites over the occipital cortex (Fig. 1B) at 2–4 s interstimulus intervals in the stimulation condition (see **Methods**), while no stimulation was applied in the no-stimulation condition. We assessed the effects of TMS on the rhythmic foot movement by quantifying the variability of the movement frequency with the SD of the cycle duration (Fig. 1A, see **Methods**). We did not observe any immediate motor effects of TMS, such as muscle twitching or movement stops. The main target area in this study was the early visual cortex. Thus, in the main text, we focused on stimulation site #11 (see Fig. 1B), which corresponds to the putative V1/V2 (pV1/V2; see Supplementary Information). Supplementary Figure S1 reports the average SD over all stimulation sites, covering broader visual areas.

Initially, we confirmed that all groups of participants successfully performed the instructed movement. The average movement frequencies over all trials in the stimulation and no-stimulation conditions were similar (0.94 ± 0.17 Hz, 1.00 ± 0.24 Hz and 0.97 ± 0.16 Hz for the acquired blind, congenitally blind and sighted groups, respectively; F_2,32_ = 0.82, p = 0.45, η_p_^2^ = 0.05).

However, we observed distinct group differences in the effect of TMS on movement performance. SD of the cycle duration in stimulation-to-pV1/V2 condition was higher when compared with no stimulation condition in the acquired blind group, resulting in the positive value of ΔSD (Increases in the SD from the no-stim condition to the stimulation-to-pV1/V2 condition; blue box plot in Fig 2A). This increase in the SD was not observed in the congenitally blind (green) or sighted groups (red). Wilcoxon signed-rank test revealed a significant increase of SD in the acquired blind group (p = 0.028 with Bonferroni correction for the three tests, *W* = 64, *r* = 0.821), but not in the congenitally blind group (p = 0.320, *W* = -28, *r* = -0.424) or the sighted group (p = 0.622, *W* = 4, *r* = 0.051) (Fig. 2A and Table 2).

**Fig. 2.**
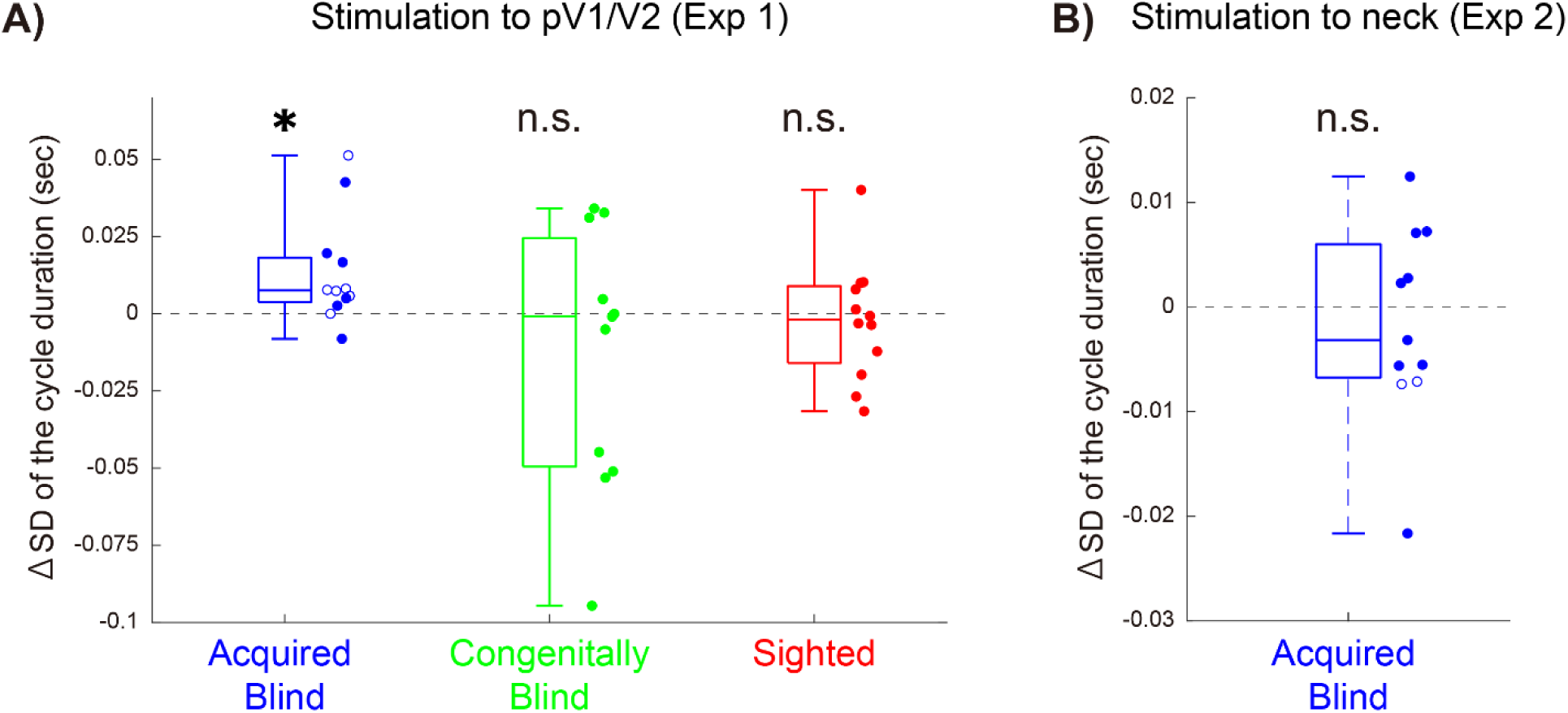
A) Experiment 1: Increases in the SD from the no-stim condition to the stimulation-to-pV1/V2 condition in which TMS was applied to stimulation site #11 for acquired blind (blue), congenitally blind (green), and sighted (red) participants. Box plots show the median and interquartile ranges of the increase in the SD for each group. Each dot represents data from an individual participant. For the blind group, open dots and closed dots indicate the subgroups of athletes (ABL^A^) and nonathletes (ABL^NA^), respectively. The asterisk indicates p < 0.05 with Bonfferoni correction. B) Experiment 2: Increases in the SD from the no-stim condition to the stimulation-to-neck condition in which TMS was applied over the right spelenius capitis (see **Methods**) for acquired blind participants. Box plots show the median and interquartile ranges of the increase in the SD for each group. Each dot represents data from an individual participant. For the blind group, open dots and closed dots indicate the subgroups of athletes (ABL^A^) and nonathletes (ABL^NA^), respectively.

**Table 2.** SD values in no-stimulation condition and stimulation-to-pV1/V2 condition and the increase in SD for each group in Experiment 1.

|  | SD<sub>no-stim</sub> | SD<sub>pV1/V2</sub> | Δ SD |
| --- | --- | --- | --- |
| **ABL (n = 12)** | **0.072 ± 0.045** | **0.085 ± 0.043** | **0.013 ± 0.017** |
| ABL<sup>A</sup> (n = 6) | 0.049 ± 0.013 | 0.063 ± 0.026 | 0.013 ± 0.019 |
| ABL<sup>NA</sup> (n = 6) | 0.095 ± 0.055 | 0.108 ± 0.045 | 0.013 ± 0.018 |
| **CBL (n = 11)** | **0.118 ± 0.077** | **0.104 ± 0.049** | **-0.013 ± 0.042** |
| **SI (n = 12)** | **0.106 ± 0.048** | **0.104 ± 0.043** | **-0.002 ± 0.019** |

*ABL, acquired blind; CBL, congenitally blind; SI, sighted. Superscripts A and NA indicate athlete and nonathlete blind participants, respectively. ΔSD = SD<sub>pV1/V2</sub> – SD<sub>no-stim</sub>. All data are reported as mean ± standard deviation across participants. The units are seconds.*

In the acquired blind group, the increase in SD with TMS was not correlated with their age (Speaman’s correlation, r = -0.432, p = 0.160), age at the onset of visual loss (r = - 0.233, p = 0.467), or duration of visual loss (r = -0.245, p = 0.442). Moreover, this increase in SD did not differ between the blind athlete (ABL^A^ in Table 2) and blind nonathlete subgroups (ABL^NA^; Mann-Whitney U test, p = 0.937 without correction, *U* = 17, *r* = 0.056 ). Furthermore, the increase in SD was not dependent on the presence or absence of light sensitivity (p = 0.202 without correction, *U* = 14.5, *r* = -0.486, Fig. S2A).

Finally, we compared the average SD over all stimulation sites with SD in the no-stimulation condition for each group. Again, we found a significant increase in the acquired blind group (p = 0.020 with Bonferroni correction for the three tests, *W* = 62, *r* = 0.795), but not in the congenitally blind group (p = 0.520, *W* = -26, *r* = 0.333) or the sighted group (p = 0.677, *W* = -26, *r* = -0.333) (Fig. S1 and Table S2).

### Experiment-2

One may argue that the increase in the SD in the acquired blind group of Experiment-1 can be caused by the sensory side effects of TMS, such as tactile sensations or a clicking sound (Duecker and Sack, 2015). To ensure that this was not the case, we conducted Experiment-2 by applying TMS to a neck muscle, the right spelenius capitis, which induces tactile/auditory sensations similar to those accompanying TMS to pV1/V2, without affecting any neural processing. We stimulated the neck muscle while 11 acquired-blind participants performed the same foot movement task as in Experiment-1 (stimulation-to-neck condition). The TMS intensity was adjusted for each participant to feel the same level of tactile sensations as those induced by TMS to the pV1/V2 (see Method). The variability in movement frequency under stimulation-to-neck conditions was compared to that in the absence of stimulation (no-stimulation condition).

The average movement frequency over all trials in the stimulation-to-neck and no-stimulation conditions was 1.012 ± 0.122 Hz, suggesting that the participants successfully performed the instructed movement. Importantly, SD of the cycle duration in stimulation-to-neck condition (0.087 ± 0.021) was almost identical to SD in no-stimulation condition (0.088 ± 0.022) and there was no significant increase in the SD from the no-stim condition to the stimulation-to-neck condition (Fig. 2B, Wilcoxon signed-rank test, p = 0.638, *W* = -20, *r* = -0.303).

## Discussion

### Overview of findings

Experiment 1 demonstrated that TMS to pV1/V2 disrupts a foot movement task in acquired blind individuals but not in congenitally blind or sighted individuals. Experiment 2 demonstrated that TMS to the neck of acquired blind individuals did not affect the movement task, suggesting that the sensory (tactile/auditory) side effects of TMS alone could not explain the disruptive effect observed in Experiment 1. Thus, our results provide the direct neurobehavioral evidence that the visual cortex of blind individuals contributes to sensorimotor control. However, this reorganization may require some visual experience before visual loss. Numerous studies have reported that reorganization in the visual cortex of blind individuals participates in nonvisual perception and cognitive functions (*2, 22, 34*). Our finding extends the knowledge on the reorganization of the visual cortex in blind individuals. Specifically, the visual cortex can be reorganized not only for perceptual and cognitive functions but also for motor function.

### Potential confounders

First, we shall discuss possible confounders of our results. One may argue that the disruptive effect of TMS (i.e., the increase in SD in Experiment 1) was attributable to advanced foot motor control rather than to acquired blindness, because six out of 11 participants were experts in the sensorimotor control of foot movements (i.e., current or former members of the Japanese national blind football team, ABL^A^). In fact, the foot movement in the ABL^A^ was more stable than the other groups, including the ABL^NA^, congenitally blind, or sighted groups, both under no-stimulation and stimulation-to-pV1/V2 conditions (Table 2). This was also observed when TMS was applied to the other stimulation sites (Table S2), indicating the strong effect of the sports experience on movement variability; this could probably be attributed to their intensive (blind football) exercises, which required controlling their foot much more accurately than in daily life.

Importantly, despite the different variability in each condition, a disruptive TMS effect (ΔSD in Table 2) was similarly observed both in the ABL^A^ and in the ABL^NA^ subgroups. Therefore, the disruptive effect of TMS observed in the acquired blind participants can not be explained by the inclusion of athletes, whose SD was lower in general. Furthermore, the movement variability in the no-stimulation condition did not account for individual differences in the disruptive effect in the congenitally blind and sighted groups (see Supplementary Information).

Another concern is that the disruptive TMS effect observed in the acquired blind group of Experiment 1 may be attributed to sensory side effects of TMS, such as tactile (pain) sensations or a clicking sound (*35*). However, Experiment 2 demonstrated it unlikely. In Experiment 2, neck stimulation did not induce disruptive effects in acquired blind participants, although the intensity of the tactile sensations and the volume of the clicking sound were similar to those induced by TMS to pV1/V2. (see Methods). Thus, our results suggest that the disruptive effect observed in Experiment 1 is primarily attributed to the neural stimulation on the brain function rather than the sensory side effects from TMS. Therefore, we reject the possible confounding factors and argue that the pV1/V2 of the acquired blind participants engaged in the foot movement.

This neck stimulation may serve as a novel and useful control stimulus during motor tasks, particularly for continuous movements. Typically, in motor TMS studies, the visual cortex is often chosen as control stimulation sites because it has been considered minimally involved in motor execution. However, recent findings suggest that the visual cortex in sighted individuals may contribute to motor control (*33*), and that plasticity mechanisms in motor and visual areas may interact (*36*), indicating that visual cortex stimulation may not always be appropriate as a control condition. Furthermore, although sham stimulation is effective for discrete movements, rapidly completed movements, it poses challenges for continuous motor tasks such as those used in the present study. Participants may experience prediction errors in the absence of expected tactile sensations, which can lead them to perceive issues with the experimental procedure and, in some cases, discontinue the task. Therefore, neck stimulation with carefully controlled levels of pain and tactile intensity could provide a novel and effective approach for assessing sensory side effects induced by TMS.

### Effects of visual experience

The disruptive effect was observed in all acquired blind participants (Fig. S2B), including five early-blind (age < 6 years), three intermediate-blind (6 ≤ age ≤ 16 years), and four late-blind participants (age > 16 years), with the exception of a late blind (#8). We did not find a significant correlation between the SD increase and age at onset of blindness. Therefore, the amount of visual experience did not appear to affect the degree of visual cortex engagement. This is consistent with the previous studies reporting the lack of relationship between amount of the visual experience and perception/cognitive functions (*20, 27, 28, 37, 38*).

In contrast to the acquired blind participants, the congenital blind participants did not show any clear disruptive effect of TMS to pV1/V2 (Fig. 2A, Table 2, and S1), suggesting that the visual experience is necessary for the engagement of the visual cortex in sensorimotor control. Visually guided movements appear early in human development, at around 6–8 months of age (*39–41*). This suggests that the neural connections between the visual cortex and sensorimotor regions, crucial for visually-guided motor control (*33*), are formed very early during development. Recent work in mice (*42*), as well as humans (*43, 44*), suggests that normal visuomotor experiences are necessary for the typical development of this connection, even although a part of the connection can be inherent (*45*). In our acquired blind participants, such neural connection for visuomotor control may be (at least somewhat) normally established at least at a very early stage of development. Then, following visual loss, these existing neural connections may be enhanced and reorganized (*1, 2, 46–48*) for sensorimotor control of nonvisually-guided movements in acquired blinds’ brain. On the other hand, such neural connections can not be developed in the congenitally blind individuals without any visuomotor experience. These might be the reason why we observed the disruptive effect of TMS to pV1/V2 on the rhythmic feet movement in the acquried blind participants, irrespective of their age at onset of blindness, but not congenitally blind participants.

### Functional roles of the visual cortex

Although the current study could not identify the exact functional role of the acquired blind participants’ visual cortex in our motor task, we can hypothesize possible roles based on the computational understanding of sensorimotor control (*49*). First, the visual cortex almost certainly does not contribute to the generation of motor commands, since TMS did not induce immediate motor effects such as muscle twitching or movement stop. Instead, the increased movement variability observed may indicate that TMS affects higher-order aspects of sensorimotor control such as temporal control processing or feedback control processing. Specifically, we speculate that TMS affected the online estimation of the body state (i.e., position, velocity, or movement phase of the foot), which is transformed into motor commands through motor regions (*49–52*). This state estimation is thought to be achieved by combining bottom-up sensory feedback and top-down sensory prediction–predicting the sensory consequences of motor commands (*53–55*). In the brain of sighted individuals, the visual cortex may contribute to the online estimation in the visual domain. Recent studies in both sighted mice and humans support the role of the visual cortex in predicting the visual consequences of movements (e.g., optical flow) through top-down connections from motor regions to the visual cortex (*42, 56–58*). This prediction is combined with visual feedback to achieve an online estimation of the body state. Similarly, in acquired blind individuals, the visual cortex may contribute to the state estimation by combining a nonvisual feedback signal and sensory prediction. This is supported by two lines of evidence: 1) acquired blind individuals maintain or even enhance connectivity between the visual cortex and sensorimotor regions following visual loss (*46–48, 59*) and 2) the visual cortex in blind individuals processes nonvisual sensory information (*1, 2*). Therefore, TMS to pV1/V2 may disrupt the online estimation of the body state, increasing movement variability in acquired blind individuals.

### Methodological limitation and future directions

There is a methodological limitation in our study. We determined the stimulation sites (see Methods) without using a magnetic resonance imaging (MRI)-guided navigation system (*17*). Thus, between-participant differences may exist in the anatomical location of stimulation sites. However, we focused on V1/V2; its stimulation site (#11) was determined based on a well-established external anatomical landmark (2.5 cm above inion (*60–63*)). Further, we performed a post-hoc simulation analysis based on MRI anatomical data. This simulation supported that our TMS procedure could prominently stimulate the V1/V2 (see Supplementary Information). Therefore, we are confident that we stimulated V1/V2 when applying TMS to stimulation site #11. On the other hand, the disruptive TMS effect may not be restricted to early visual cortices. The analysis of the mean TMS effect across all sites (Fig. S1, Table S1) showed a significant effect in which TMS increased the movement variability only in the acquired blind participants. Further research is needed to examine the possibility that a broader neuronal network, including higher-order visual cortices, reorganizes to contribute to sensorimotor control in blind individuals.

## Conclusion

Our study provide the direct neurobehavioral evidence that the visual cortex of blind individuals contributes to sensorimotor control. Our findings indicate that the human brain’s plasticity following sensory loss is more flexible than previously thought, i.e., functional repurposing of the lower sensory cortices is not restricted to perception and cognitive functions but also extends to motor function. This plasticity may increase the neural resources available for sensorimotor control in blind individuals, helping them interact with the everchanging and diverse environment.

## Methods

### Participants

Eighteen acquired blind (i.e. lost vision after birth, #1–#12, #37–#42 in Table 1 and 3), 12 congenitally blind (i.e. no visual experience, #13–#24) and 12 sighted participants (#25–#36) participated in either or both of the two experiments: Experiment-1 and Experiment-2. All participants were healthy volunteers without a history of cognitive impairment or psychiatric disorders. Congenital blindness was defined as total blindness (vision limited to light perception or worse) diagnosed before one year of age, whereas acquired blindness was defined as the same condition diagnosed at or after one year of age. All congenitally blind participants reported that they had never experienced color or form vision since birth. They presented no contraindications to TMS, which was assessed using a screening questionnaire in compliance with the guidelines for noninvasive magnetic brain stimulation in research applications (*64*). The causes of blindness for all blind participants were related to lesions in the prechiasmatic visual system. These experiments were approved by the ethics committee of the National Institute of Information and Communications Technology and conducted according to the Declaration of Helsinki. All participants provided written informed consent before participating in the experiment and were naïve to the purpose of the study.

**Table 3.**
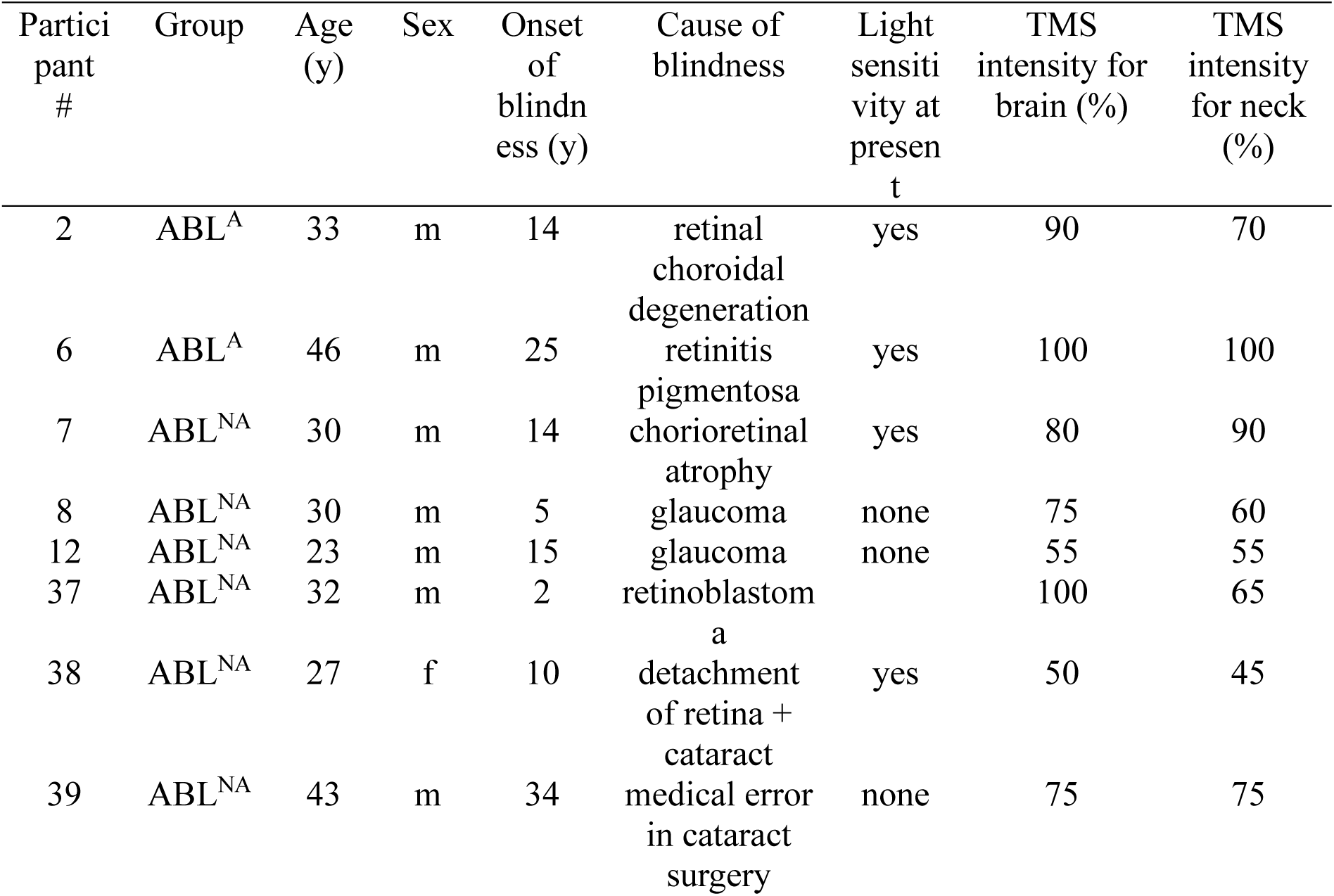
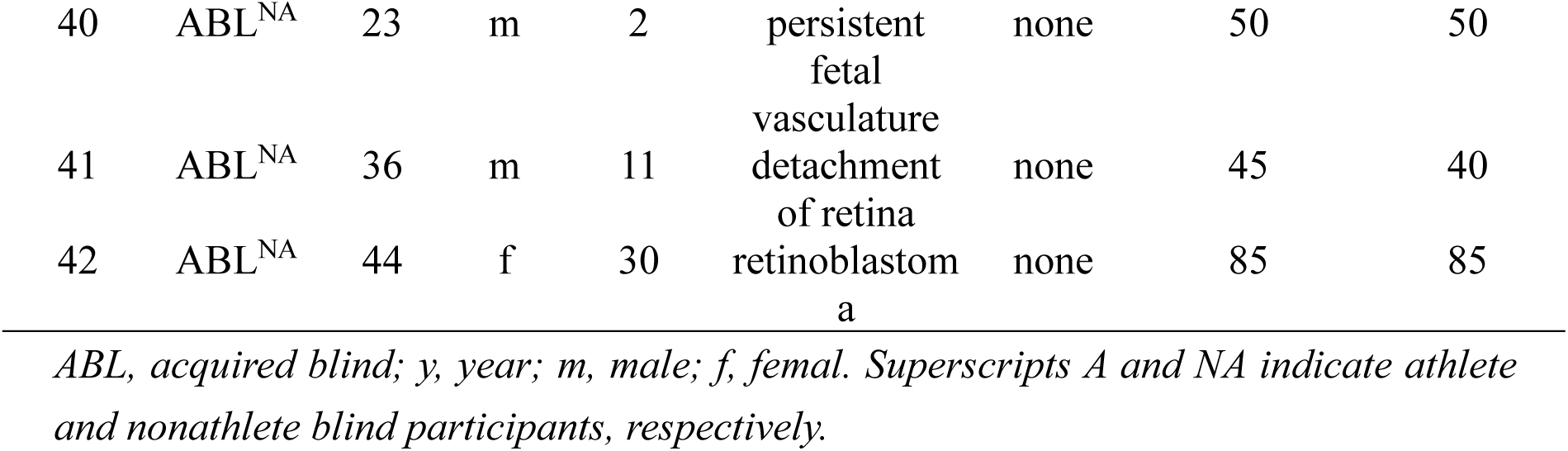
Characteristics of the participants in Experiment 2.

**Table 4.** SD values in no-stimulation condition (no-stim) and stimulation-to-neck condition and the increase in SD in Experiment 2.

|  | **SD<sub>no-stim</sub>** | **SD<sub>neck</sub>** | **ΔSD** |
| --- | --- | --- | --- |
| **ABL (n = 11)** | **0.088 ± 0.022** | **0.087 ± 0.021** | **-0.002 ± 0.009** |

*ABL, acquired blind. ΔSD = SD<sub>neck</sub> − SD<sub>no-stim</sub>. All data are reported as mean ± standard deviation across participants. The units are seconds.*

#### Experiment 1

Twelve acquired blind (acqureid blind group: ABL in Table 1), 12 congenitally blind (congenitally blind group: CBL) and 12 sighted (sighed group: SI) individuals with normal or corrected-to-normal vision participated in Experiment-1. Age was similar between the acquired blind participants (age 27.50 ± 8.37, mean ± standard deviation (SD) across participants) and sighed participants (27.75 ± 7.24), while both were younger significantly than the congenitally blind participants (41.00 ± 7.01; F_2,32_ = 11.74, p = 1.50×10^-4^, η_p_^2^ = 0.42). The acquired blind group comprised two subgroups: athlete (ABL^A^) and nonathlete (ABL^NA^). The six blind participants in the ABL^A^ subgroup were current or former members of the Japanese national blind football team. The remaining six in the ABL^NA^ subgroup were blind individuals from the general population.

One congenitally blind participant (#12) was former members of the Japanese national blind football team, while the other participants in the CBL were from the general population. All sighted participants belonged to the general population. We determined the sample size (n = 12 per group) based on the effect of TMS on motor tasks assessed by previous studies (*65–68*).

#### Experiment 2

Eleven acquired blind individuals (see Table 3), including two athletes (#2, #6) and three nonatheletes (#7, #8, #12), who participated in Experiment 1, and six nonathletes who did not participate in Experiment-1 (#37–#42), took part in Experiment 2. We determined a sample size (n = 11) similar to Experiment-1.

### Apparatus

#### Experiment 1

All participants wore eye patches and an eye mask to eliminate any possible visual stimuli ∼30 min before the experiment and continued to wear them throughout the experiment. Furthermore, they were instructed to close their eyes during the experiment. This was to ensure that the sighted participants performed the task without any visual information as did blind participants. They sat in a comfortable reclining chair with their heads on a chinrest and legs extended on a leg rest. The height and location of the chin and leg rests were adjusted for each participant to comfortably make the rhythmic movements of alternating dorsiflexion and plantar flexion of their feet. An electromagnetic position sensor (Micro Sensor 1.8 Extra Flex, Polhemus Liberty, Burlington, VT) recorded the six dimensional kinematic data (position: x, y, and z; orientation: azimuth, elevation, and roll) of the first metatarsals of both feet. The obtained data were digitized with a temporal sampling ratio of 240 Hz and then low-pass filtered at a cutoff frequency of 5 Hz.

Electromyographic (EMG) activity was recorded at 2,000 Hz utilizing surface electrodes from the medial and lateral heads of the gastrocnemius and tibialis anterior of both legs using wireless EMG sensors (Wave Plus Wireless EMG, Cometa, Bareggio, Italy). This EMG recording was prepared to capture possible muscle twitches with TMS. However, since we did not observe any twitches, EMG data were not reported in this study.

TMS stimulation was applied using a 70-mm figure-eight coil and a Magstim Rapid Transcranial Magnetic Stimulator (Magstim Company, Spring Gardens, UK). The maximum stimulator output was 2.1 T. TMS pulse delivery was controlled by an in-house program running on MATLAB, version R2013b (The MathWorks, Natick, MA).

#### Experiment 2

The experimental apparatus was the same as Experiment1 except for the following two points. First, as all participants were blind, they neither wore eye patches nor eye masks. Second, EMG activity was not measured.

### TMS protocol

#### Experiment 1

Fourteen stimulation sites were arranged in a 2 × 7 grid and numbered from left to right and top to bottom (Fig. 1B); these sites were identified for each participant using an elastic swimming cap with 15 markers indicating the inion position and the 14 stimulation sites. The marker positions were determined using a cap worn by a male individual with an average head size for Japanese men (head circumference: ∼57 cm (*69*)) who did not participate in the experiment. The markers were placed at inion, 2.5 cm (stimulation site #11), and 5 cm (stimulation site #4) dorsal from the inion on the median line (nasion-to-inion line through the vertex), and 2, 4, and 6 cm right and left away from the median line at the same level as stimulation sites #4 and #11. Each participant wore the swimming cap on the head with 1) the inion marker placed at the participant’s inion, 2) stimulation sites #4 and #11 aligned on the participant’s median line, and 3) stimulation site #11 placed at 2.5 cm dorsal from the inion. Thus, stimulation site #11, the main target in this study, was well controlled among participants (on the median line and 2.5 cm dorsal from the inion). On the other hand, the other stimulation sites could not be strictly controlled.

Subsequently, the stimulation intensity for each participant was determined using the highest value that did not induce scalp pain (Table 1). Single TMS pulses were applied to each stimulation site (sites #1–#14 in order). For stimulation site #1, stimulations were applied by gradually increasing the intensity by 5% starting from 60% intensity of the maximum stimulator output until the participant reported any pain. Then, the intensity was decreased by 5% until the pain disappeared. The last intensity was defined as tentative intensity. When the participant did not report pain with a stimulation at 100% intensity, the tentative intensity was set to 100%. For other stimulation sites, we began stimulation with the tentative intensity. When the participant reported pain, we decreased it by 5% until pain disappeared and updated the tentative intensity. This procedure was repeated for all sites up to site #14.

The TMS intensity used in the experiment was comparable between the acquired blind (77.08% ± 11.97% of the maximum stimulator output), congenital (72.73% ± 16.18%), and sighted (84.58% ± 9.64%) groups (F_2,32_ = 2.56, p = 0.09, η_p_^2^ = 0.14).

#### Experiment 2

TMS was applied to a neck muscle. The stimulation site was on the right spelenius capitis, ∼2 cm superior from the horizontal line passing through the 7^th^ spinal process. The TMS intensity for the neck (I_Neck_) was adjusted for each participant to feel the same level of tactile sensations as induced by TMS to pV1/V2 (stimulation site #11). I_Neck_ determination consisted of four steps. In STEP 1, we determined the intensity of the stimulation to pV1/V2 as in Experiment-1 (I_Brain_). Then, we provided three single-pulse stimulations to pV1/V2 with an inter-pulse interval of ∼1s and asked the participant to remember the tactile sensation induced by TMS to pV1/V2 with I_Brain_. In STEP 2, we applied TMS to the neck muscle to determine I_Neck_. We gradually increased the intensity by 5% starting from 40% of the maximum stimulator output until the participant felt a tactile sensation comparable to that induced by TMS to pV1/V2. The last intensity was defined as I_Neck_. In STEP 3, to confirm that the tactile sensations induced by these two stimulations were comparable, we again provided one single-pulse stimulation to pV1/V2 with I_Brain_ followed by one single-pulse stimulation to the neck with I_Neck_. When the participants confirmed that the sensations were comparable, the determination procedure was completed. Otherwise, we moved to STEP4; when reporting that the tactile sensation of the neck stimulation was stronger or weaker, we decreased or increased I_Neck_ by 5% until reaching a comparable level of tactile sensation to that induced by TMS of the pV1/V2 with I_Brain_. Then, we went back to STEP 3 (confirmation process). STEP 4 and STEP 3 were performed until the participants confirmed that the sensations were comparable.

The TMS intensity of the pV1/V2 was similar between participants in Experiment 2 (I_Brain_: 73.18% ± 20.28%) and the acquired blind group in Experiment 1 (77.083% ± 11.968%). Slightly weaker stimulations induced the same level of tactile sensations when applying TMS to the neck muscle (I_Neck_: 66.82% ± 19.27%). The loudness (sound pressure level, SPL) of the clicking sound induced by TMS to the neck (69.79 ± 0.44 dB; mean ± SD of ten measurements) was comparable to that induced by TMS to pV1/V2 (72.67 ± 0.84 dB). SPL was measured by a sensor placed around a participant’s right ear while receiving TMS to the neck with average intensity in Experiment 2 (67%) or to pV1/V2 with average intensity in Experiment-1 (77% for the blind group).

### Task procedure

#### Experiment 1

The participants were blindfolded and while sitting on the chair, made rhythmic alternative movements of dorsiflexion and plantar flexion of the feet without any sound cue. To maintain the movement frequency as constant as possible, the participants practiced the movements with a sound frequency of 1 Hz for ∼2 min twice: once before the determination process of the stimulation intensity and once before starting the experiment. They were instructed to maintain the consistent movement frequency throughout the experiment. No instructions were provided regarding the motion range of the foot movement.

In each experimental trial, the rhythmic movement was performed under each of the following three conditions. During the stimulation condition, 20 single TMS pulses were applied to one of the 14 stimulation sites at interstimulus intervals of 2–4 s (drawn from a uniform random distribution). The TMS coil was held tangentially to the scalp. The trial was started 3 s before the first stimulation and ended 3 s after the last. Each trial lasted 56.34–70.26 s. During the sham condition, a fake coil was placed over stimulation site #11, while another coil delivered 20 single TMS pulses in the air near the fake coil. Thus, the participant could feel the clicking sound coming from the fake coil on the scalp. The duration and timing of TMS pulses were determined according to the protocol used in the stimulation condition. During the no-stimulation condition, no coil was placed on the scalp, and TMS pulses were not generated. The trial duration was set at 50 s.

The participants completed two blocks of 15 trials, including 14 stimulation trials (one for each of the 14 stimulation sites) and one sham trial. The trial order was pseudorandomized for each block. One no-stimulation trial was performed before the first block and after the second block. Thus, the participants completed a total of 32 experimental trials.

#### Experiment 2

The procedure of Experiment-2 was similar to Experiment 1. The participants underwent the same ∼2 min practice with a 1 Hz sound, before and after the determination process of the stimulation intensity. The participants were instructed to maintain the foot movement frequency at 1 Hz throughout the experiment without specific instructions on the motion range. Then, the participants completed two blocks of 10 trials each. For each trial, they performed the same task as Experiment-1. In each block, five odd-numbered trials were in the stimulation-to-neck condition, and five even-numbered trials were in the no-stimulation condition. In total, the participants performed 10 stimulation trials and 10 no-stimulation trials in total.

In the stimulation condition, the TMS pulse was applied to the neck muscle (the right spelenius capitis). Interstimulus interval, number of single TMS pulse, and trial’s start and end points were identical to the stimulation condition in Experiment 1. As the neck is concave, we could not adhere the coil to the skin surface of the stimulation site. Thus, we held the coil so that the center position was 2–3 cm away from the skin surface. During the no-stimulation condition, no coil was placed on the neck, and the TMS pulses were not generated. The trial duration was set at 50 s, as for the no-stimulation condition in Experiment 1.

### Data analysis

#### Experiment 1

The effects of TMS on movement were examined by evaluating movement variability. For each foot in each trial, we initially identified the duration of each movement cycle, defined as the time between the adjacent dorsiflexion peaks (Fig. 1A and see details in the next paragraph), and calculated the SD of the durations from all movement cycles. Then, we obtained the SD for each stimulation site by averaging the four SD values for left and right foot in the two stimulation trials (first and second blocks). Similarly, the SD in the no-stimulation trials was calculated as control. Our main analysis focused on stimulation site #11, pV1/V2 (see TMS protocol). The other stimulation sites might have covered broad visual areas, including early (V1/V2) and higher-order (V3, V4, and V5) visual areas, according to the literature (*61, 63, 70, 71*). To observe the overall trend of the TMS effect on early and higher-order visual cortices, we analyzed the average SDs of all stimulation sites as shown in Fig. S1.

The dorsiflexion peaks (Fig. 1A) for each foot in each trial were identified as follows. First, the time average of the elevation angle from the horizontal plane was calculated and subtracted from the entire time series to obtain the time series of a relative angle (*θ* in Fig. 1A). Then, the crossing times where *θ* changed from negative to positive were identified. Finally, the point of maximum value between each of two consecutive crossing times was identified as a peak. The identification code is available online (see Data availability).

#### Experiment 2

First, the SD was calculated in each trial as in Experiment 1. Then, for each stimulation-to-neck and no-stimulation condition, all SD values (10 trials for each condition) were averaged to obtain the SD.

### Data exclusion

Data from the sham trials were not analyzed since almost all participants noticed the lack of stimulation and/or felt something going wrong during the trials. Hence, the data were probably biased by this surprising effect.

#### Experiment 1

Participant #17 was excluded from the analysis. In the middle of the 1st block (after 9^th^ trial), the participant reoprted a discomfort with the position of the feet and head and changed the position as well as removed the chin rest. Thus, we analyzed data from 11 participants for the congenitally blind group. For stimulation site #14 of participant #35 (Table 1) and sites #2, #4, #8, #13 of participants #19, data from only the second block were utilized to calculate the SD owing to the lack of the first block data due to a recording error. For stimulation site #5 of participant #20, the second block data of the left foot were excluded as the foot posture changed significantly during the trial. For stimulation sites #4 and #7 of participant #21, the second block data of the left foot were excluded as he fell asleep during the trial. In total, 0.82% of the stimulation trials were excluded from analysis, while all the no-stimulation trials were analyzed.

#### Experiment 2

For the left foot (participant #2, 8, 12, 37, 40, and 42) or both feet (#7, 38, and 39) of some participants, the azimuth angle was used, which is the angular orientation of the foot in the horizontal projection plane. This was because we had observed, upon visual inspection, that their azimuth angle reflected the dorsiflexion peaks more clearly than the elevation angle, given their internally-rotated leg posture. For participant #2 (Table 3), data from only the second block (5 trials for each condition) were utilized to calculate the SD because of the lack of the first block data due to a recording error.

### Statistical analysis

Because of the violation of normality, the SD was examined by the two-tailed Wilcoxon signed rank test or the two-tailed Mann-Whitney U test for within- and between-participant comparisons, respectively. Nonparametric Spearman’s rank correlations were used to assess the relationship of the SD with the following variables: age, age at the onset of visual loss, or duration of visual loss. For the other data analysis, we analyzed parametric one-way ANOVA for the three-group comparisons or two-tailed unpaired t-tests for two-group comparisons. The Bonferroni correction was used for multiple comparisons. All analyses were conducted using MATLAB R2024a (The MathWorks, Natick, MA). The significance level was set at 0.05 in all the analyses. For the Wilcoxon signed rank test, the *p* value, the test statistic *W*, and the effect size r were reported (*72*). For the Mann-Whitney U test, the *p* value, the test statistic *U*, and the effect size r were reported (*72*). We reported effect size with partial eta squared (η_p_^2^) for the ANOVAs, Cohen’s d_z_ for the paired *t*-test, and Cohen’s d for the unpaired *t*-test (*73*). All data are presented as mean ± SD across participants.

#### Experiment 1

To examine the effects of TMS on stimulation site # 11 (pV1/V2), the Wilcoxon signed-rank test was used to test diffences in the SD between the no-stimulation and stimulation conditions for each group. The same Wilcoxon signed-rank test was conducted for the average SD of all stimulation sites (Fig. S1). Because the acquired blind group showed a significant increase in SD on pV1/V2, the Spearman’s rank correlation analyses were performed to examine the relationship of the SD increase ( Δ SD in Table 2; SD in stimulation-to-pV1/V2 condition minus SD in no-stimulation condition) with age, age at onset of visual loss, and duration of visual loss. Furthermore, to evaluate the relationship between SD increase and the participants’ sports experience, the ABL^A^ and ABL^NA^ subgroups were compared using the Mann-Whitney U test. To examine whether the presence of light sensitivity affects SD increase, two subgroups of acquired blind participants with and without light sensitivity were compared using the Spearman’s rankSpearman’s rank (Fig. S2A). To examine differences in age, TMS intensity, and movement frequency between groups, a one-way ANOVA was performed. When a significant main effect was observed, the effect of group was examined using a unpaired t-test.

#### Experiment 2

The effects of TMS to the neck muscle were compared by the SD in the no-stimulation condition and the stimulation-to-neck condition using the Wilcoxon signed rank.

## Data availability

Data (time of the dorsiflexion peak for all participants and a sample foot movement trajectory) and codes to reproduce Fig. 2 and S1 and the related analyses are available from https://github.com/ikegami244/Blind-TMS. The code to identify the dorsiflexion peaks is also available from the same repository.

Code to replicate the simulation reported in the Supplementary Information is available at https://github.com/satoshi-hirose/Blind-TMS.TMS-simulation-with-simNIBS. The anatomical MRI data are not publicly available due to ethical restrictions but are available from the corresponding author on reasonable request.

## Acknowledgments

We thank Dr. Kaoru Amano and Dr. Nobuhiro Hagura for their valuable comments on the first draft of this manuscript, Dr. Mitsuaki Takemi for his help in the TMS simulation analysis, and Dr. Keisuke Tani for his suggestion in the neck stimulation. We thank Ms. Naoko Katagiri, Dr. Kenta Yamamoto, Ms. Noriko Karasudani, Ms. Mayura Fujita, and Ms. Keiko Chiba for their help in conducting the experiments and organizing the data. We thank Mr. Shigeo Murakami and Mr. Hajime Teranishi of the Japanese Blind Football Association for his help in recruiting participants. TI was supported by JSPS KAKENHI Grant #16K12974 and JST, PRESTO #JPMJPR22S1. EN was supported by JSPS KAKENHI Grant #JP19H05723.

## SUPPLEMENTARY INFORMATION

### TMS simulation to identify the stimulation site

We conducted a simulation analysis to investigate whether the TMS used in Experiment 1 could stimulate the early visual cortices (V1 and/or V2). Specifically, we used anatomical MRI data from each participant to simulate the electrical field induced in the visual cortex by TMS at stimulation site #11.

#### Participants

Fifteen (eight acquired blind and seven sighted) out of the 24 individuals from the acquired and sighted groups who participated in Experiment 1 participated in this simulation study. The rest of the participants were excluded because of contraindications to MRI (one sighted and four blind) or video recording problems in the TMS experiment (four sighted).

#### Procedures

We acquired T1-weighted anatomical brain images of the participants (MP-RAGE; repetition time [TR]: 1900 ms; echo time [TE]: 2.48 ms; inversion time [TI]: 900 ms; flip angle: 9°, field-of-view [FOV]: 256 mm × 256 mm; voxel size: 1.0 mm × 1.0 mm × 1.0 mm) with a 3.0-T SIEMENS scanner (Trio Tim, with a 32-channel head coil). Then, we performed the following simulation analysis for each participant to identify the electrical field induced by TMS to stimulation site #11.

#### Simulation

We conducted the simulation for two trials on stimulation site #11 for each participant, following the procedure introduced by Saturnino et al. (2019). We first segmented the MRI volume into the different head tissues, and the tissue surfaces were identified. Then, we simulated the electrical field induced by stimulation by the TMS coil placed at 2.5 cm dorsal from the inion (stimulation site #11). From the simulated electrical field, the intensity of the electrical field (norm of the electrical field vector; V/m) in the cortical gray matter was mapped on the cortical surface (Fig. S3).

The simulation for each participant requires three parameters: the location and orientation of the TMS coil and the stimulus intensity. For the location, we first identified the inion (Iz), the nasion (Nz), and the vertex (Cz) on the scalp surface based on the morphometrical structure of the head surface. Then, we identified the point 2.5 cm above Iz on the circle passing through Iz, Nz, and Cz. For the orientation, we identified the TMS coil orientation in the actual TMS experiment by visual inspection of the video recording of the experiment by one of the authors (TI). The coil orientation in each of the two trials for stimulation site #11 was identified in 5° increments. The stimulus intensity used in the simulation was identical to that used in the TMS experiment. We used the default values for the other simulation parameters, including the tissues’ electrical conductance.

We used the headreco pipeline (Nielsen et al., 2018) with Statistical Parametric Mapping (SPM; https://www.fil.ion.ucl.ac.uk/spm/software/spm12) and Computational Anatomy Toolbox for SPM (https://neuro-jena.github.io/cat) with default parameters for the segmentation. We performed the simulation with simNIBS software (simNIBS 3.2; https://simnibs.github.io/simnibs). The anatomical identification of the cortical area was based on the human cortical parcellation provided by the Human Connectome Project (HCP-MMP1.0; Glasser et al., 2016).

## Results

We performed two analyses with the simulation data. First, we examined whether the intensity peak of the simulated electrical field was observed in V1/V2 (Table S3). The peak of the electrical field was observed within V1 or V2 in 14 out of 15 participants (Table S3) in both trials. An exception was observed in Participant #2, whose peak was in V3A (7.4 mm apart from the edge of V1).

Next, we checked whether the intensity of the electrical field induced in V1/V2 was sufficient to affect neural processing in these regions. We regarded an electrical field > 10 V/m as sufficient to affect neural processing. The threshold was set according to previous studies showing that neurons fire in response to an artificially induced electrical field with ∼ 10 V/m intensity (Park et al., 2013, Bonmassar et al., 2012, Chan and Nicholson 1986). The proportion of the V1/V2 surface area where the induced electrical field was above the threshold (stimulated region) is 81% ± 10% (mean ± standard deviation; see Table S3).

These two results suggest that the TMS procedure used in Experiment-1 substantially stimulated V1/V2, with sufficient stimulus intensity.

### The movement variability in the no-stimulation condition did not account for individual differences in the disruptive effect in the congenitally blind and sighted groups

To deny the possibility that the disruptive effect of TMS (i.e., the increase in SD in Experiment 1) was attributable to advanced foot motor control rather than to acquired blindness, we have shown that both athletes (ABL^A^) and non-athletes (ABL^NA^) in the acquired blind group exhibited disruptive TMS effects (i.e., increases in SD). To further strengthen this argument, we evaluated the relationship between the original motor performance, assessed using SD in the non-stimulation condition, and the TMS effect.

Data from 23 participants in the congenitally blind and sighted groups were included in this analysis. Participants were divided into two subgroups—high SD (N = 12) and low SD (N = 11)—based on their SD values in the first no-stimulation condition conducted at the beginning of the experiment. The TMS effect was assessed by subtracting the SD of the second non-stimulation condition conducted at the end of the experiment, from the average SD of the two TMS stimulation-to-V1/V2 conditions. By using statistically independent values for subgroup assignment (SD of the first non-stimulation condition) and for the calculation of the dependent variable (SD of the second non-stimulation condition), we avoided the statistical bias known as regression to the mean.

As a result, no significant TMS effect was observed in either the high-SD (t(11) = -0.80, p = 0.44) or low-SD (t(10) = 0.06, p = 0.96) subgroups. Furthermore, no evident relationship was observed between SD in the initial non-stimulation condition (x-axis in Figure S4) and the TMS effect (y-axis).

## Supplementary figures

**Fig. S1.**
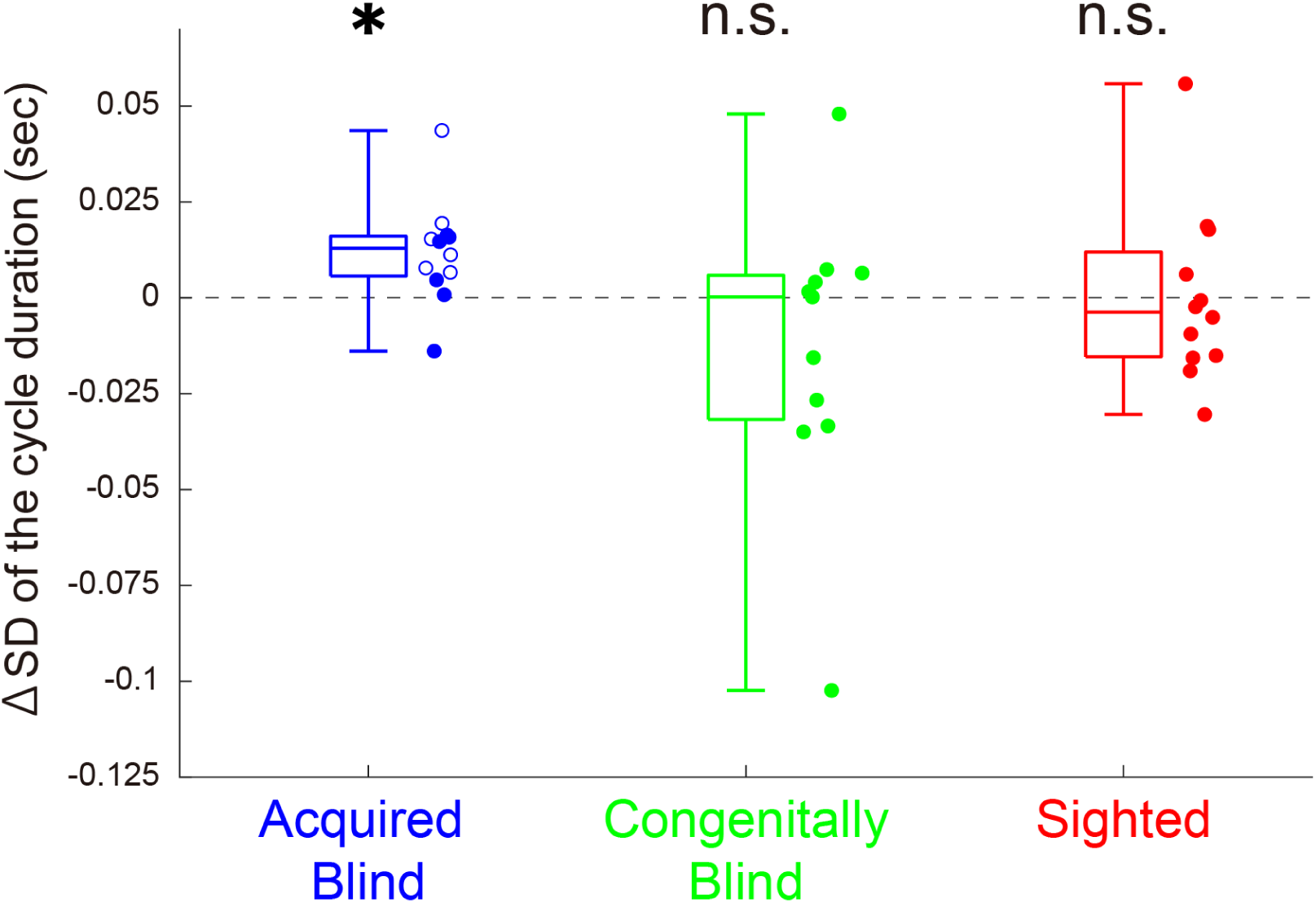
The average SD over all stimulation sites (all sites) was compared with the SD in the no-stimulation condition. The increases from the SD of the no-stim condition to the average SD over all stimulation sites for acquired blind (blue), congenitally blind (green), and sighted (red) groups in Experiment-1. Box plots show the median and interquartile ranges of the increase in the SD for each group. Each dot represents data from an individual participant. For the blind group, open dots and closed dots indicate the subgroups of athletes (ABL^A^) and nonathletes (ABL^NA^), respectively. The asterisk indicates p < 0.05 with Bonfferoni correction.

**Fig. S2.**
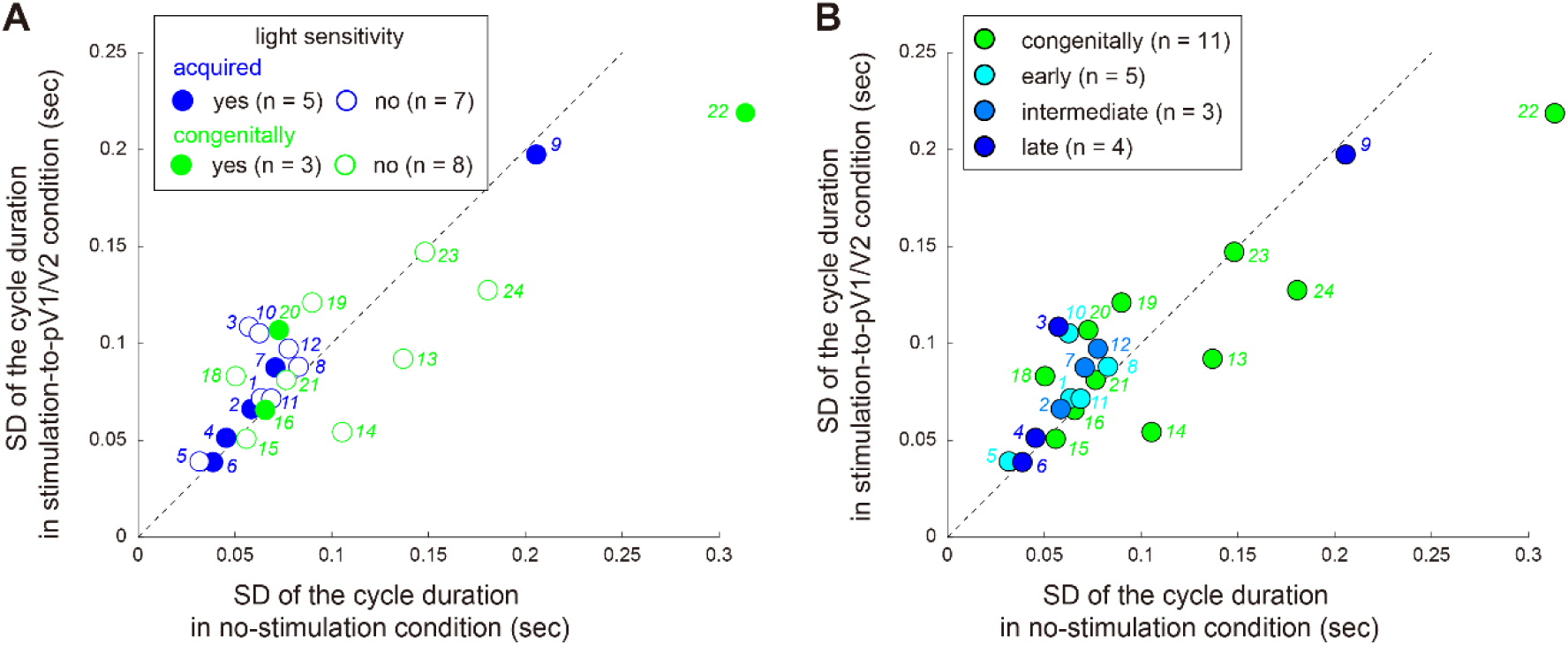
SD of the cycle duration for the blind participants in the no-stimulation condition (x-axis) and the stimulation-to-pV1/V2 condition (y-axis) of Experiment 1. A) For each group of acquired (blue) and congenitally (green) blind participants, two subgroups with and without light sensitivity (filled and open circles, respectivly) were compared. The dashed line indicates y = x. In the acquired blind participants, TMS to pV1/V2 increased the SD in the cycle duration relative to the no-stimulation condition in all participants exept for one (#8, see Table 1). The increase in SD did not differ between the two subgroups. In contrast, no clear trend was not observed among the congenitally blind participants nor between the twosubgroups. B) Early-, intermediate-, late-acquired blind participants (light, intermediate, and dark blue circles, respectively), and congenitally blind participants (green circle) are compared. The dashed line indicates y = x. No clear difference was observed among the three subgroups of the acquired blind participants.

**Fig. S3.**
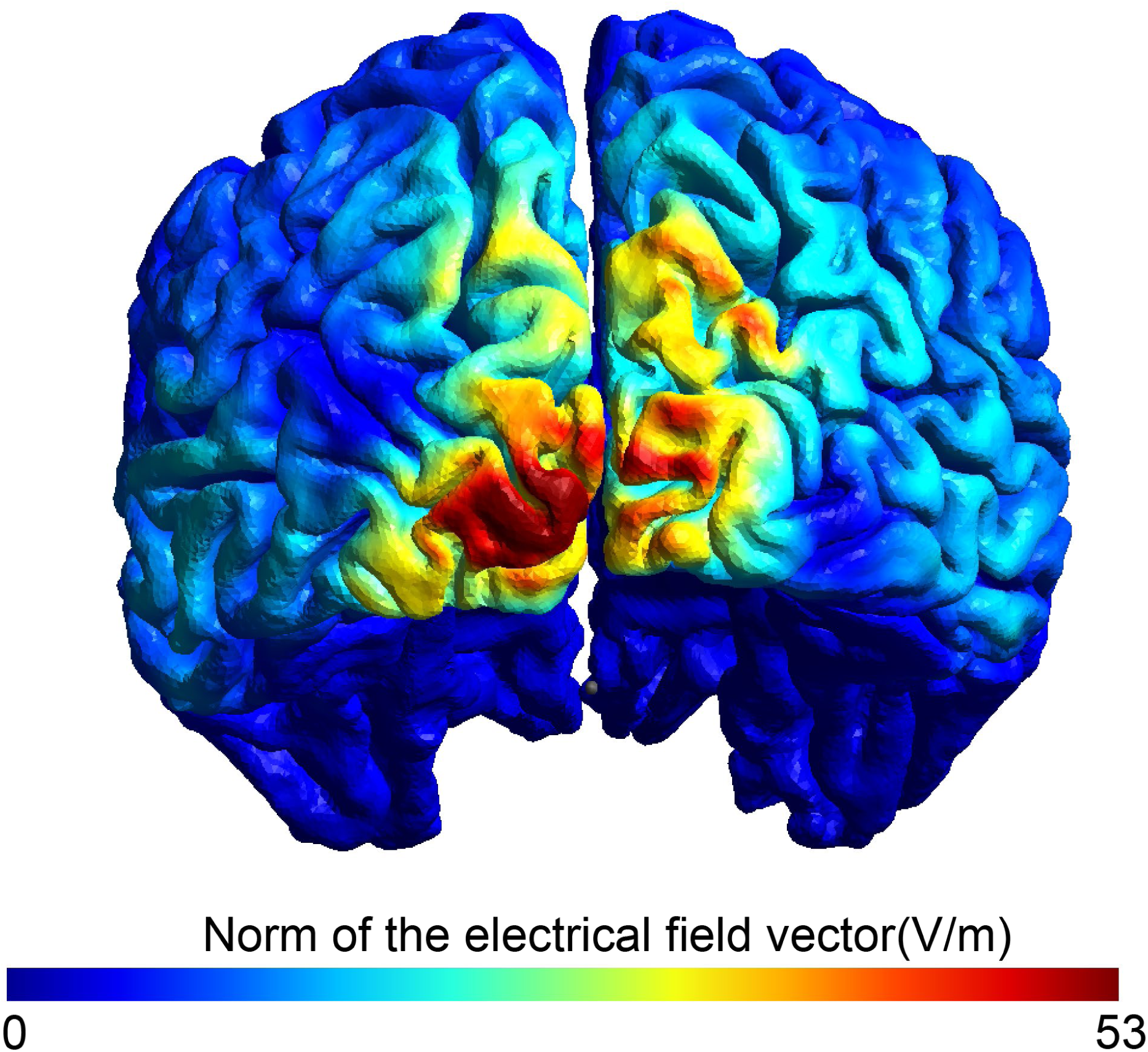
Distribution of the electrical field intensity (norm of the electrical field vector; V/m) in the cortical gray matter in a typical participant (participant #6). The intensity in the cortical gray matter was mapped onto the cortical surface. The figure shows a posterior view of the cortex.

**Fig. S4.**
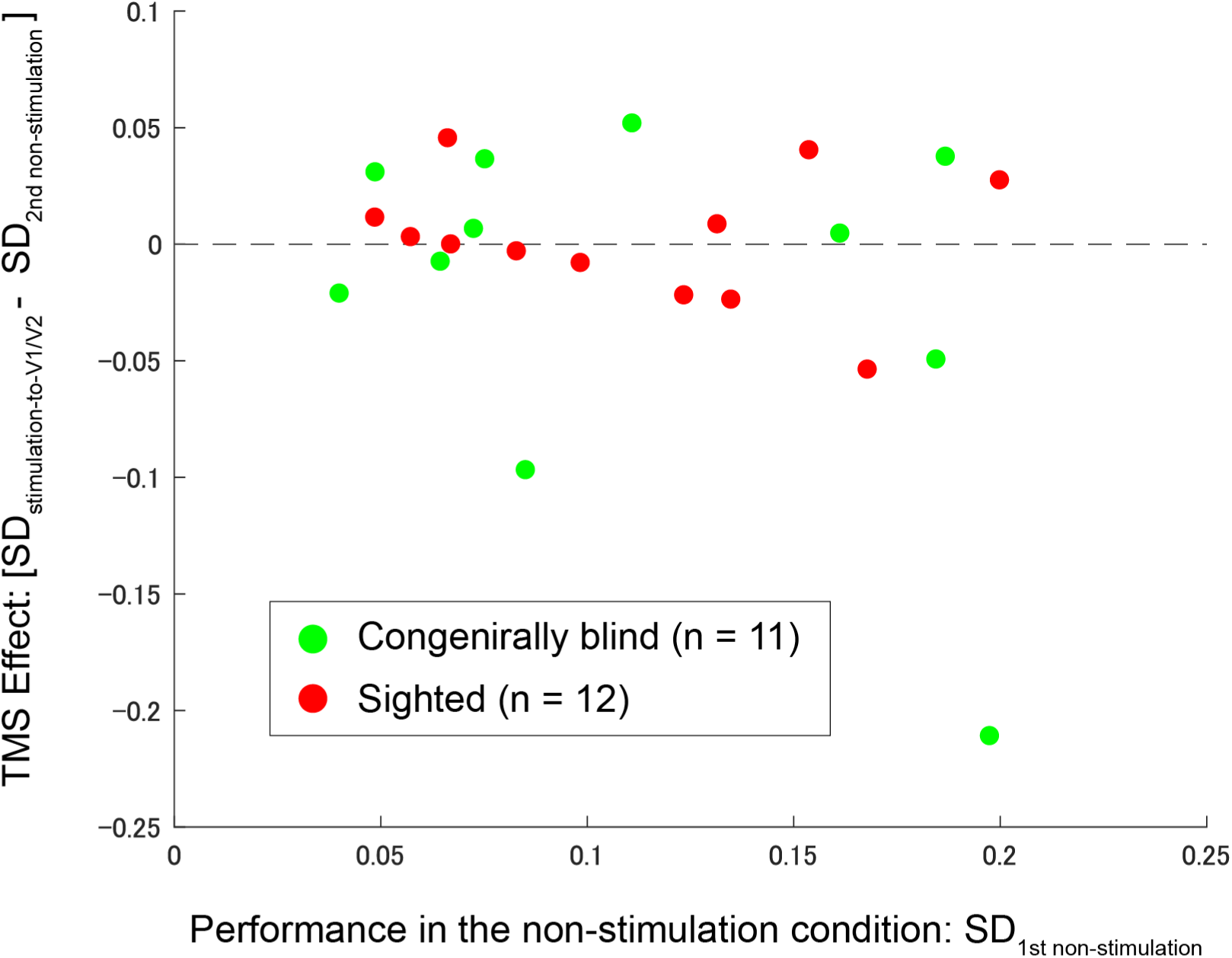
Relationship between performance in the non-stimulation condition and TMS effects: The performance in the non-stimulation condition was assessed using the SD from the first non-stimulation condition. TMS effects were evaluated by subtracting the SD of the second non-stimulation condition from the average of the two sets of TMS stimulation-to-V1/V2 conditions.

**Table S1.**
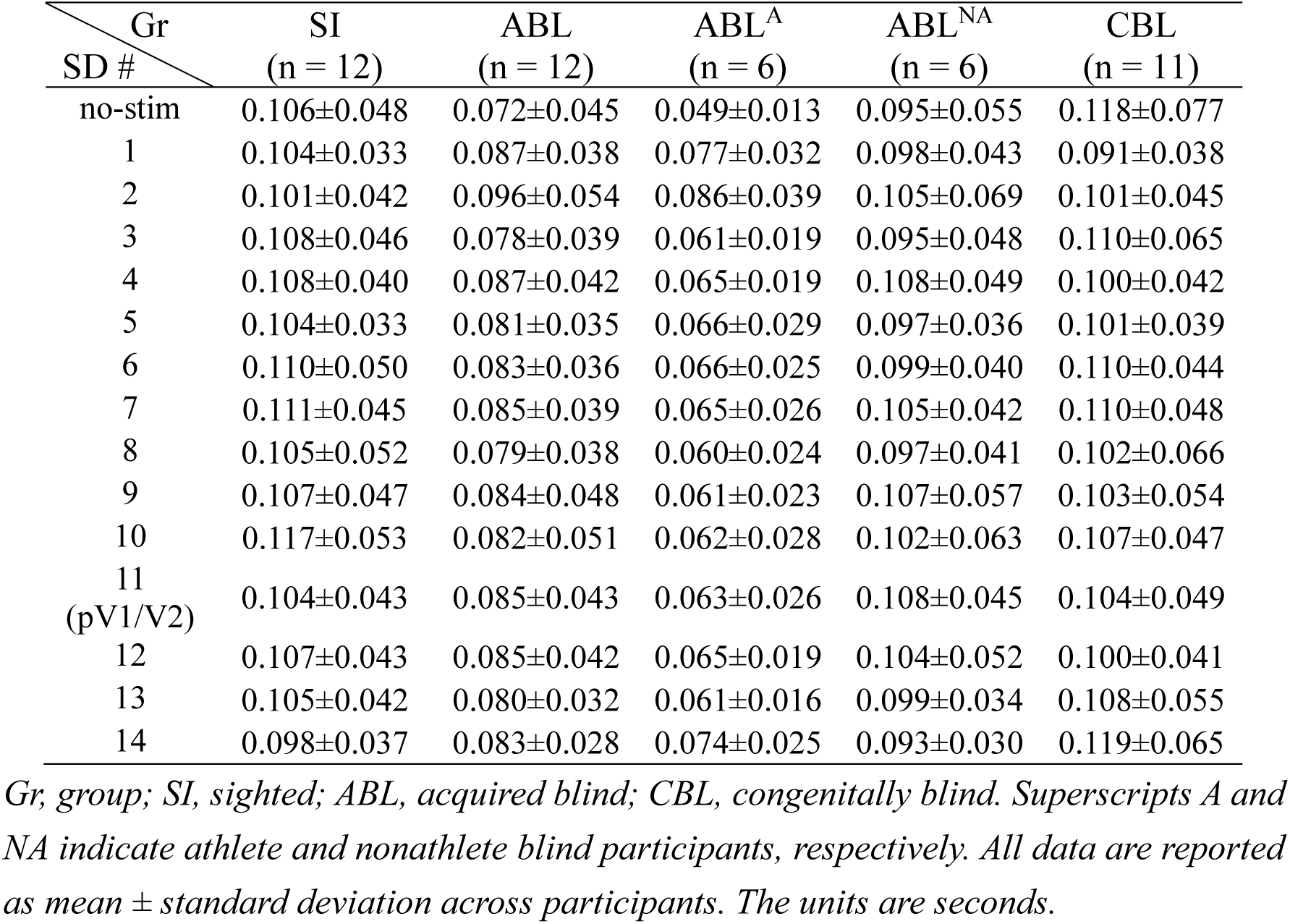
SD of the cycle duration in all stimulation sites (#1-14) and the no-stimulation condition (no-stim) for the participants of Experiment-1 and the congenitally blind participants.

**Table S2.** Simulation results.

| Participant<br># | Group | Peak location | proportion of stimulated region<br>in V1/V2 (%) |
| --- | --- | --- | --- |
| 1 | ABL<sup>A</sup> | Left V1/Right V1* | 88.5 |
| 2 | ABL<sup>A</sup> | Left V3A | 54.9 |
| 4 | ABL<sup>A</sup> | Left V1 | 81.3 |
| 6 | ABL<sup>A</sup> | Left V1 | 80.7 |
| 7 | ABL<sup>NA</sup> | Left V1 | 76.5 |
| 8 | ABL<sup>NA</sup> | Left V1 | 83.5 |
| 10 | ABL<sup>NA</sup> | Right V1 | 91.4 |
| 11 | ABL<sup>NA</sup> | Left V2 | 88.7 |
| 28 | SI | Left V2/Right V1* | 86.6 |
| 30 | SI | Right V1 | 86.8 |
| 31 | SI | Left V2 | 77.7 |
| 33 | SI | Left V1 | 81.0 |
| 34 | SI | Left V1 | 87.8 |
| 35 | SI | Left V1/Left V2* | 90.3 |
| 36 | SI | Left V1 | 80.8 |

\* Asterisks indicate that the peaks for the first and second trials were located in different anatomical areas, where results were indicated as follows: first trial peak location / second trial peak location.

